# Continuous and widespread population interactions in the Jōmon society via geometric morphometrics on 3D data of human crania

**DOI:** 10.1101/2025.02.09.637294

**Authors:** Hisashi Nakao, Akihiro Kaneda, Kohei Tamura, Koji Noshita, Mayu Yoshida, Tomomi Nakagawa

**Affiliations:** Department of Anthropology and Philosophy, Nanzan University, Japan; Nara National Research Institute for Cultural Properties, Japan; The Center for Northeast Asian Studies, Tohoku University, Japan; Graduate School of Environmental Studies, Tohoku University, Japan; Center for Integrated Japanese Studies, Tohoku University, Japan; Graduate School of Science, Kyushu University, Japan; Graduate School of Letters, Nagoya University, Japan

**Keywords:** Jōmon, human skeletal remains, geometric morphometrics, three-dimensional (3D) data, resilience

## Abstract

This study examines the morphological variations within and between regional populations via geometric morphometrics of larger samples of three-dimensional data of Jōmon human crania (*N* = 363, including 146 females, 215 males and two unknown-sex individuals) from 97 sites to investigate the population interactions in the period, which Japanese anthropologists and archaeologists have extensively discussed. The results show that morphological variations are more pronounced within individual populations especially in the principal component one, relating to the facial width, degree of prognathism, and location of occipital areas, in contrast to the relatively smaller variations observed between different phases and geographical regions. This observation is consistent with the possibility that the population interactions of the Jōmon people had been widespread and continuous, and which has an important implication for their resilience against severe climate changes at that time: The relative stability of the Jōmon society might be sustained by their frequent interactions with various populations, as suggested by insights from relevant archaeological, ethnographic, and genetic research.

## Introduction

Anthropologists have extensively discussed the population history in the Jōmon period of the Japanese archipelago (14,000 to 800 cal BC, incipient: 14,000 to 9,300 cal BC, early (including initial in this study): 9,300 to 3,400 cal BC, middle: 3,400 to 2,400 cal BC, late: 2,400 to 1,200 cal BC, final: 1,200 to 800 cal BC (Taniguchi 2019), although the incipient period was excluded in the present research because no human skeletal remains are found in this period), when people engaged mainly in hunting and gathering. Some research on human skeletal remains including crania and limbs has supported the notion of relative uniformity within the Jōmon population, which could be sustained by widespread population interactions, although others have claimed that spatiotemporal differences may exist to some extent (Dodo 1982, 1986; Hanihara & Ishida 2009; Kondo 1994, 2018; Mouri 1988; Ogata 1981; Yamaguchi 1982a, 1982b). In particular, it has often been claimed that geographical clines are found from north to south because the Jōmon people was assumed to be influenced by both the southern and northern populations outside the Japanese archipelago. Due to the dietary conditions, Ogata (1981) also argued that the body size of the incipient and early Jōmon people was smaller than that of the other subperiods.

Many archaeological studies have also discussed population interactions via a widespread distribution and exchange of artifacts such as pottery and lithics over extensive geographic regions during this period (Abe 2010; Fukunaga 2020; Habu 2004; Imamura 1996, 2011; Taniguchi 2019; Yano 2016). It has often been pointed out that a certain type of pottery was widely distributed across regions. An example is the Ento-kaso type distributed in both the Hokuriku and the northern Tohoku regions of the early phase. Even though the distribution of a type of pottery was regionally restricted, it could be affected by a distant region. The Funamoto type of the middle phase was distributed mainly in the Sanyo regions, and this type of pottery strongly influenced the pottery in the Kyushu region in the same phase. In the late phase, when samples were obtained from the most various regions in the present study, it was found that a large amount of the Horinouchi type I pottery, originating from the Kanto region was excavated in the Kinki region, and that the Nakatsu type pottery from the Kinki region was transported to the Kanto region and influenced the origin of the Shomyoji type pottery there (Imamura 2011; Fukunaga 2020; Suzuki 2011; Yano 2006). Such diverse and widespread distribution of pottery suggests that the Jōmon people interacted widely, although it has often been argued that Jōmon society was regionally diverse and distinct, including settlement styles, lifestyles, and some types of archaeological remains such as clay figurines. This is the reason why the Jōmon cultural complex has been called as ‘Jōmon cultures’ (e,g., Matsumoto 2005; Taniguchi 2019; Yamada 2019).

The present study examines the population interactions in the Jōmon society in an anthropological way, i.e., through geometric-morphological analysis of the cranial features of the Jōmon people, using a much larger three-dimensional (3D) cranial sample collected from broader regions than previous relevant studies. The geometric morphometrics of 3D data has gained traction in various fields, extending to objects such as lithics (García-Medrano et al. 2020; Hashemi et al. 2021; Shott et al. 2010; Yahalom-Mack et al. 2020). Although the morphological variations of the Jōmon people have been explored in various previous studies depending on the traditional biodistance method, the present study has the following novelties. 3D data of Jōmon crania have rarely been examined by geometric morphometrics with some exceptions (e.g., Buck et al. 2019, 2024; Makishima and Ogihara 2009; Nakao et al. 2023, 2024), and these exceptions still focused on smaller samples from restricted regions. Although 3D data may have richer information than traditional biodistance data, less biased larger samples are important for estimating macroscopic population interactions. Furthermore, when analyzing the resulting data, geometric morphometrics is able to track morphological variation as a whole, whereas the traditional biodistance method has to compare each measured distance independently. These advantages would provide some new insights that previous studies could not find.

The results indicate that variations within phases and regions are more salient than between them, which suggests that population interactions in the Jōmon society was continuous and widespread. Some environmental implications of the results, especially their relevance to resilience against the climate changes will be also discussed.

## Materials and Methods

We collected 3D cranial data from the Jōmon period (14,000 to 800 cal BC, incipient: 14,000 to 9,300 cal BC, early (including initial in this study): 9,300 to 3,400 cal BC, middle: 3,400 to 2,400 cal BC, late: 2,400 to 1,200 cal BC, final: 1,200 to 800 cal BC (Taniguchi 2019)) in collaboration with local archaeological centers, museums, and universities. We employed a range of scanning technologies, specifically Creaform HandySCAN BLACK, HandySCAN BLACK™ Elite, and Einscan HD, in addition to Structure from Motion and Multi-View Stereo techniques based on 2D photographs (Kaneda et al. 2022; Nakagawa et al. 2022a). Previous studies have confirmed the consistency of the models generated by the aforementioned methods for human crania (Nakagwa et al. 2022a; Nakao et al. 2022). Our dataset comprises 363 Jōmon crania (including 146 females, 215 males, and two individuals with unknown sex), with specific information available in Table 1 and S1 within the supplementary materials. These 3D data were collected from various regions spanning the island of Honshu within the Japanese archipelago, encompassing regions from Tohoku to Kyushu (Figure 1). To provide a basis for comparison, we included cranial data from the Kuma–Nishioda site of the Yayoi period (800 BC to 250 AD), comprising 22 crania located in northern Kyushu region.

**Table 1.** Data for the crania examined in this study; F: female, M: male, U: sex unknown.

| Area | Initial |  |  | Early |  |  | Middle |  |  | Late |  |  | Final |  |  | Unknown |  |  | Total |  |  |
| --- | --- | --- | --- | --- | --- | --- | --- | --- | --- | --- | --- | --- | --- | --- | --- | --- | --- | --- | --- | --- | --- |
|  | F | M | U | F | M | U | F | M | U | F | M | U | F | M | U | F | M | U | F | M | U |
| Tohoku |  |  |  | 1 | 1 |  | 9 | 14 |  | 6 | 14 |  | 4 | 4 |  | 6 | 5 |  | 26 | 38 | 0 |
| Kanto |  | 1 |  | 3 | 1 |  | 10 | 20 |  | 14 | 35 |  | 1 |  |  |  |  |  | 28 | 57 | 0 |
| Hokuriku | 1 | 1 |  | 2 | 6 |  |  | 2 |  |  | 1 |  |  |  |  |  |  |  | 3 | 10 | 0 |
| Tokai | 3 | 3 |  |  | 1 |  |  |  |  | 4 | 11 |  | 42 | 48 | 1 |  |  |  | 50 | 63 | 1 |
| Kinki | 2 |  |  |  |  |  |  | 1 |  | 1 |  |  |  |  |  | 1 | 1 |  | 4 | 2 | 0 |
| Sanyo |  | 1 |  |  | 1 |  | 7 | 10 |  | 4 | 3 |  | 15 | 16 |  |  | 1 |  | 26 | 31 | 0 |
| Sanin |  |  |  |  |  |  |  |  |  |  |  |  |  |  |  |  |  |  | 0 | 0 | 0 |
| Shikoku | 2 | 2 | 1 |  |  |  |  |  |  | 1 | 1 |  |  |  |  |  |  |  | 4 | 3 | 1 |
| Kyushu |  |  |  |  | 1 |  | 1 | 1 |  | 2 | 6 |  |  |  |  | 4 | 2 |  | 7 | 10 | 0 |
| Total | 8 | 8 | 1 | 6 | 11 |  | 27 | 48 |  | 32 | 71 |  | 62 | 68 | 1 | 11 | 9 |  | 146 | 215 | 2 |

**Figure 1:**
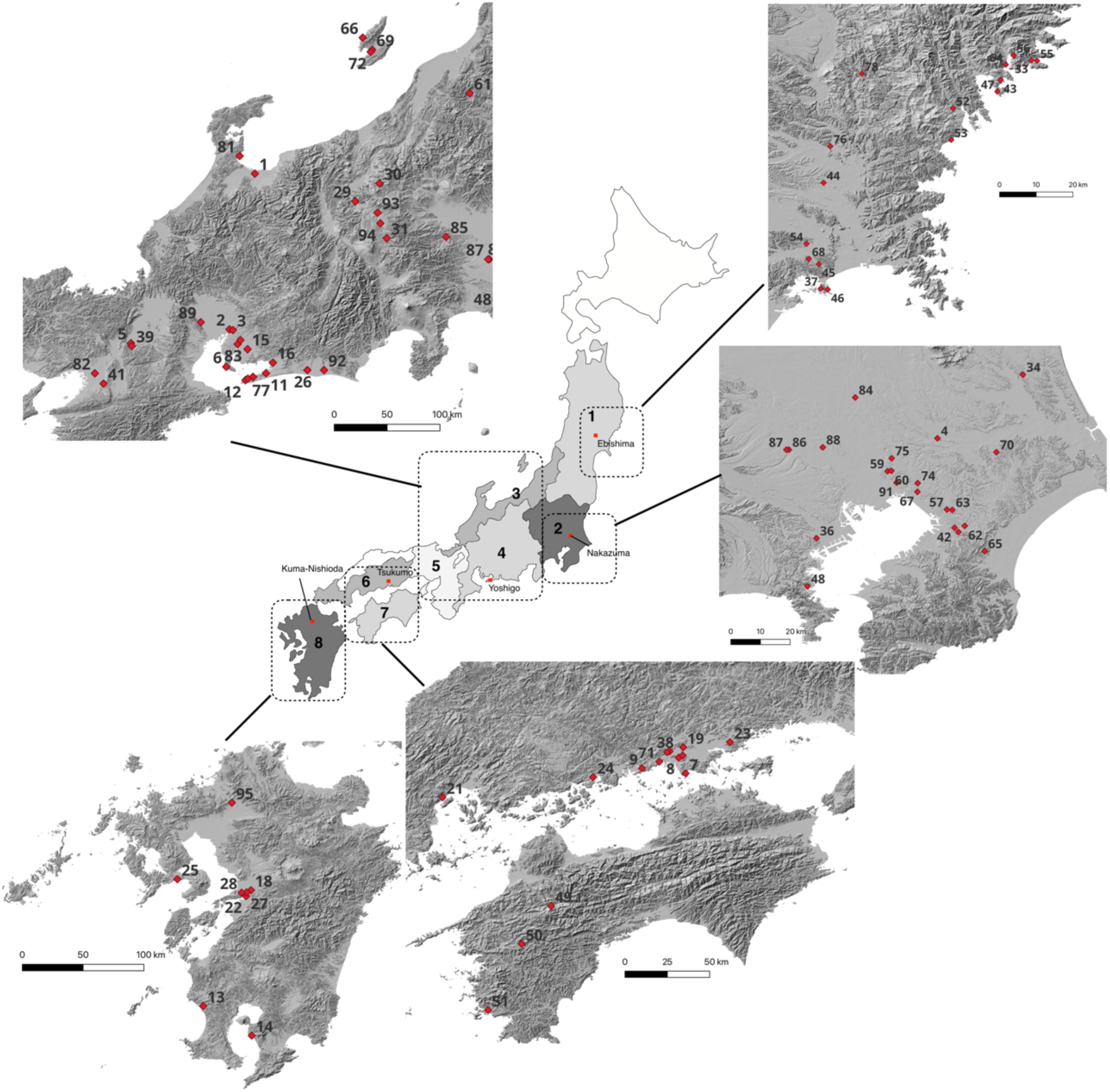
Areas divided and site locations investigated in this study: 1. Tohoku, 2. Kanto, 3. Hokuriku, 4. Tokai, 5. Kinki, 6. Sanyo, 7. Shikoku, 8. Kyushu. The map is based on the color altitude map published by the Geospatial Information Authority of Japan with information on the sea area from Hydrographic and Oceanographic Department, Japan Coast Guard, and modified by HN using QGIS (3.20.3). Site locations: 1. Odake, 2. Tamanoi, 3. Oguruwa, 4. Nakazuma, 5. Awazukotei, 6. Hayashinomine, 7. Azuhashiride, 8. Fukuda, 9. Tsukumo, 10. Ikawazu, 11. Yoshigo, 12. Kawaji, 13. Ichiki, 14. Kunugibaru, 15. Horiuchi, 16. Inariyama, 17. Harasaki, 18. Adaka, 19. Hatori, 20. Satogi, 21. Miyazono, 22. Ono, 23. Ohashi, 24. Ota, 25. Ropponmatsu, 26. Shijimizuka, 27. Sobata, 28. Todoroki, 29. Boji, 30. Yukura cave, 31. Tochibara iwakage, 32. Ebishima, 33. Miyano, 34. Wakaumi, 35. Daizennominami, 36. Shosenzuka, 37. Satohama, 38. Satoki, 39. Ishiyama, 40. Suzumimatsu, 41. Kou, 42. Ariyoshiminami, 43. Monzen, 44. Aoshima, 45. Kawakuradi hibiki, 46. Murohama, 47. Nakasawahama, 48. Shomyoji, 49. Kamikuroiwa iwakage, 50. Nakatsugawado, 51. Hirajo, 52. Tagara, 53. Maehama, 54. Higashiyogai, 55. Nonomae, 56. Ohora, 57. Kusakari, 58. Sotomeyachi, 59. Akiyamamukaiyama, 60. Nakabyo, 61. Muroya, 62. Takada, 63. Kasori north, 64. Shimofunato, 65. Shimooda, 66. Sadootsuka, 67. Fujisaki, 68. Kaigarazuka, 69. Sannomiya, 70. Soyaaranami, 71. Nakatsu, 72. Fujizuka, 73. Kumanobayashi, 74. Takanekido, 75. Kainohana, 76. Kaitori, 77. Hobi, 78. Kumaana, 79. Isodori-Ebimori, 80. Iyasakadaira, 81. Tomari cave, 82. Morinomiya, 83. Motokariya, 84. Shinmei, 85. Myoonji, 86. Mizuko, 87. Okkoshi, 88. Enshoji, 89. Hazawa, 90. Imokawa, 91. Kosaku, 92. Nishi, 93. Shimekake, 94. Getsumeisawa, 95. Kuma-Nishioda. The map is based on the content (hillshade) provided by the Geospatial Information Authority of Japan (GSI) as a part of the GSI Tiles collection, and modified by HN using QGIS (3.20.3).

The file size of the original 3D data is so large that it takes a long time to read the data and locate landmarks in R, ranging from 100 to 600 MB depending on the measurement methods. To address this issue, we reduced the size to 2.5%–15% of the original using Meshlab (https://www.meshlab.net/). Previous studies have confirmed that this reduction process does not result in a model that is significantly different from the original (Kaneda et al. 2022; Noshita et al. 2022a, 2022b).

Some prehistoric crania are incomplete, so we attempted to reconstruct lost areas in 3D models. This reconstruction involved mirroring remaining portions along the median line, passing through nasion, prosthion, and bregma (Fantini et al. 2008; Martin 1928; Nakagawa et al. 2022b; Nakao et al. 2023, 2024). In cases where mirroring was not possible, particularly when corresponding parts were entirely lost, we applied the estimate missing function, employing the thin-plate-spline method within the geomorph package (version 4.0.4) of R (version 4.2.1) and R studio (2022.07.0 + 548) (Adams et al. 2022; R Core Team 2020; RStudio Team 2020).

Sex and age estimation of skeletal remains depended mainly on published excavation reports, where anthropologists estimated the sex and age. If any relevant information was not given in the reports, we estimated them by investigating pelves and crania (especially post-cranial features and sizes of the zygomatic arch and mastoid process) for sex (Buikstra and Ubelaker 1994), and the facies symphysialis (Brooks and Suchey 1990) and cranial suture for the age (Meindl and Lovejoy 1985; Sakaue 2015).

We chose 31 representative and/or less complicated landmarks, which were placed on 3D models using the geomorph package in R (Figure 2). After conducting generalized procrustes analysis to rotate and adjust the landmark configurations for all Jōmon and Yayoi crania, we performed principal component analysis (PCA) to investigate the spatiotemporal variations of these crania. We did not perform PCA for each phase or region because no region has samples from all phases, and no phase has samples from all regions. Additionally, we also performed Steel–Dwass tests to evaluate the principal component scores of Jōmon crania based on each phase and region. All statistical analyses were performed using R studio.

**Figure 2:**
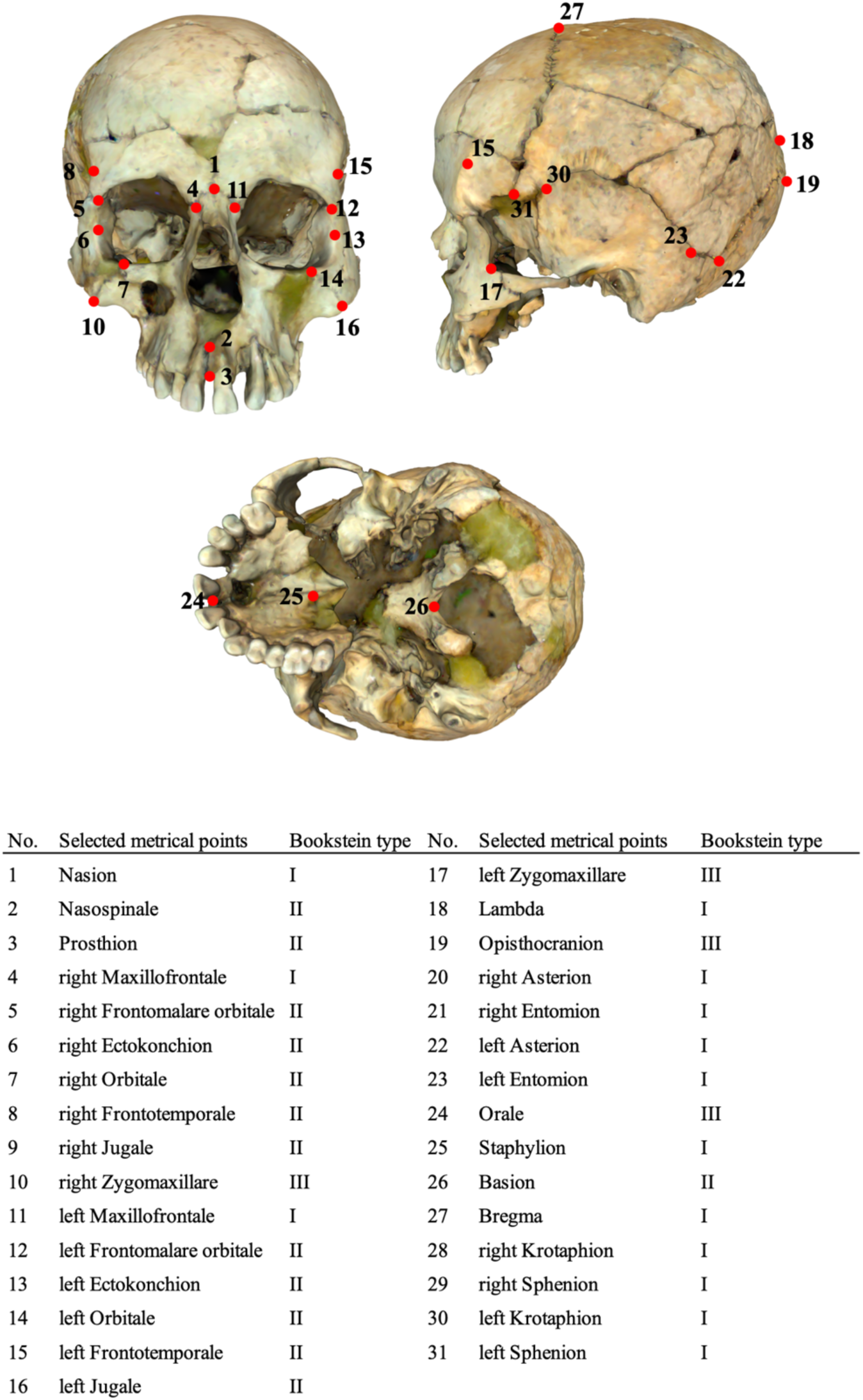
Locations of landmarks; 3D data is from the cranium of the Miyano shell midden, owned by the Ofunato City Museum; locations are selected based on traditional Martin (1928) and Caple and Stephan (2016). 9, 20, 21, 28, and 29 are not depicted.

Our research focuses on a human trait, specifically the cranium, enabling us to use the landmark method, much like similar studies (Badawi-Fayad & Cabanis 2007; Kuzminsky et al. 2108; Makishima & Ogihara 2009; Zelditch et al. 2012). Landmark selection was based on the traditional Martin (1928) and the more recent Caple and Stephan (2016) (see Figure 2 for the Bookstein type of landmark selected in the present study).

After conducting PCA on all samples of the Jōmon crania, we specifically selected cranial data from Jōmon sites with relatively larger sample sizes to compare the morphological and related genetic variation with less morphologically and genetically diverse populations from another period, i.e., the Yayoi period. These Jōmon sites include Ebishima, Nakazuma, Yoshigo, and Tsukumo, and their locations are indicated in Figure 1 and supplementary S1. We conducted a comparative analysis with the data from the Kuma–Nishioda site using geometric morphometrics and PCA. This choice is motivated by the common assumption that Yayoi people, especially in the northern Kyushu region, might exhibit lower genetic and morphological diversity due to a genetic bottleneck caused by gene flow from migrating Korean populations (Naito 1981; Nakahashi & Nagai 1989; Nakao et al. 2023, 2024). The data from Kuma–Nishioda site are therefore useful for assessing the extent of genetic and morphological diversity among Jōmon people.

## Results

The PCA results indicate cumulative contribution rates exceeding 75% up to the 20^th^ principal component (PC) (Table 2). To streamline the discussion, this study primarily focuses on PCs with magnitudes larger than 5%, PC1–PC5. PC1 positively correlates with narrow facial width, prognathism, and high and short occipital areas, while PC2 is positively linked to temporal length and negatively related with facial length. PC3 primarily captures temporal length, and PC4 reflects facial height. PC5, primarily representing cranial asymmetry, is not included in the analysis as it is influenced by ground pressure during burial (Figure 3). We also deform a cranium from the Fujizuka shell midden (ID: 341, see supplement S1) according to the morphological change each PC captured (Figure 4).

**Table 2:** Contribution rates and cumulative proportion in principal component analysis.

|  | PC1 | PC2 | PC3 | PC4 | PC5 | PC6 | PC7 | PC8 | PC9 | PC10 |
| --- | --- | --- | --- | --- | --- | --- | --- | --- | --- | --- |
| Contribution rate | 11.1% | 9.0% | 6.4% | 6.1% | 5.4% | 4.9% | 4.1% | 3.6% | 3.0% | 3.0% |
| Cumulative Proportion | 11.1% | 20.1% | 26.5% | 32.6% | 38.0% | 42.9% | 47.0% | 50.6% | 53.6% | 56.6% |
|  | PC11 | PC12 | PC13 | PC14 | PC15 | PC16 | PC17 | PC18 | PC19 | PC20 |
| Contribution rate | 2.7% | 2.5% | 2.3% | 2.1% | 2.0% | 1.8% | 1.7% | 1.6% | 1.5% | 1.4% |
| Cumulative Proportion | 59.3% | 61.8% | 64.1% | 66.2% | 68.2% | 70.1% | 71.8% | 73.4% | 74.9% | 76.4% |

**Figure 3:**
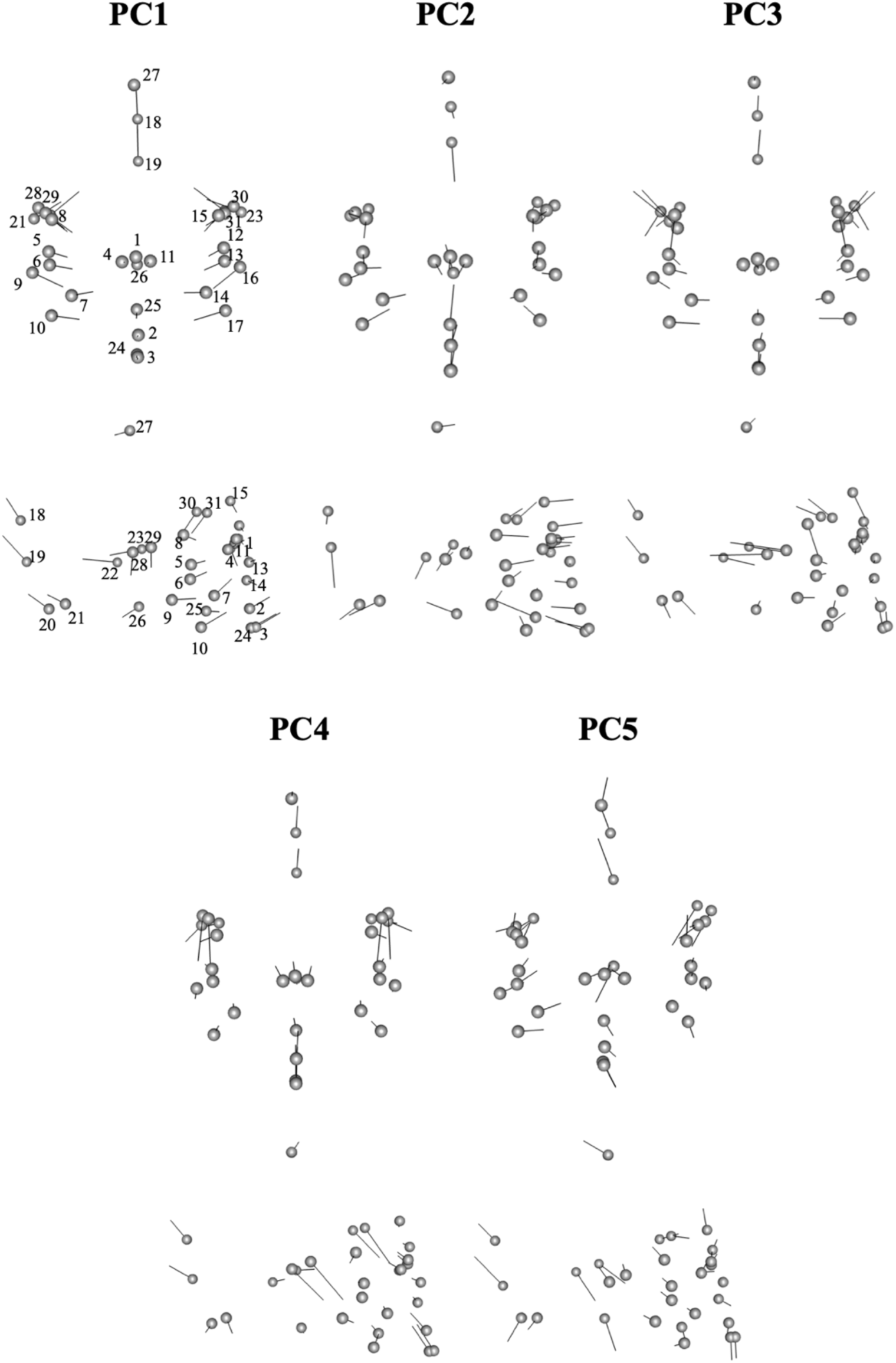
Configuration changes captured by each principal component. The numbers correspond to ones in Figure 2.

**Figure 4:**
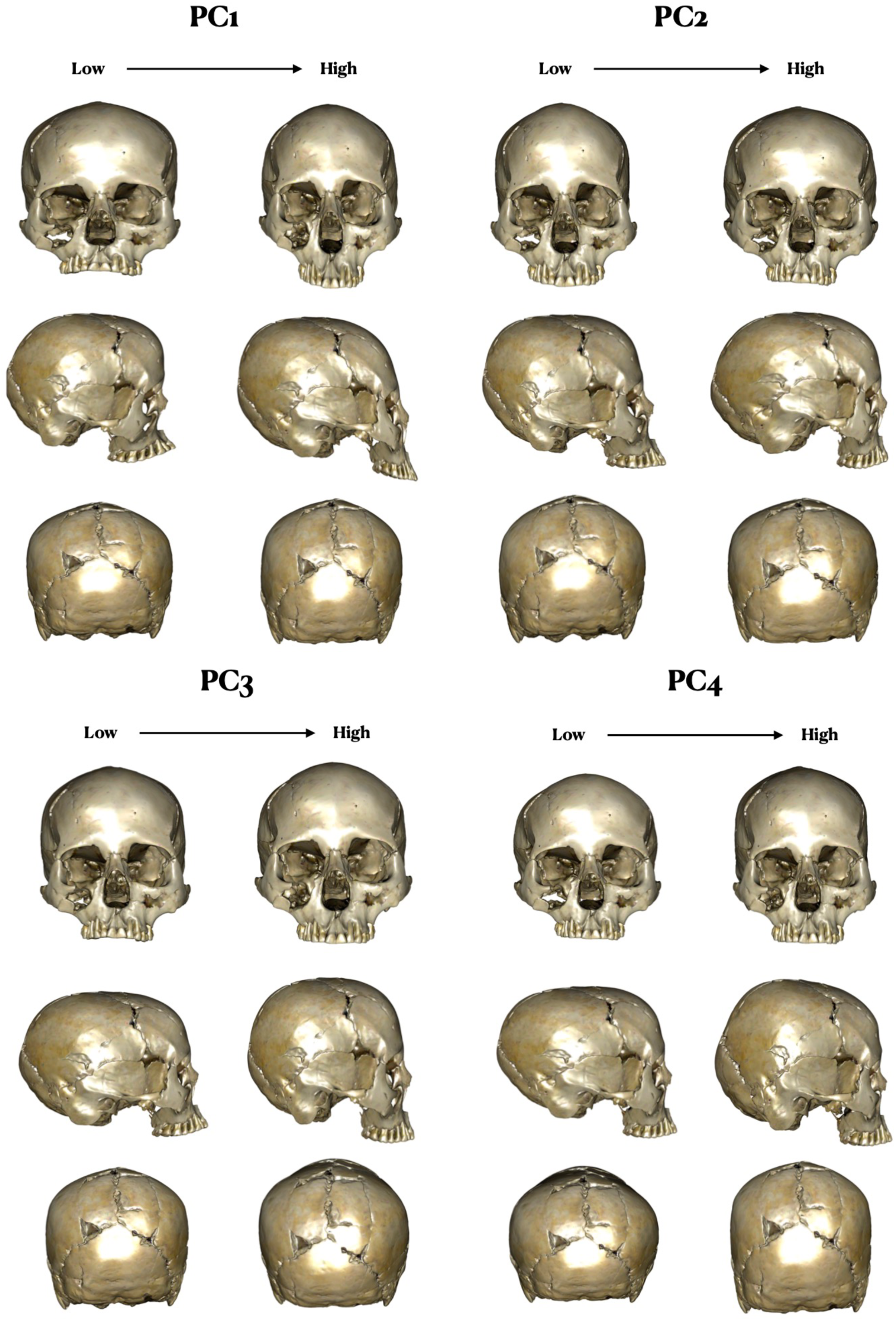
A deformed human skeletal remain (excavated from the Fujizuka shell midden) according to the morphological change of each PC.

The PCA results, graphically presented in Figures 5 and 6, suggest that the differences between phases are relatively small, in line with the statistical tests on PCs that indicate no significant differences. Notably, the scatter plots in Figure 5 and the boxplots in Figure 6 show more prominent regional differences, consistent with the statistical tests on PCs showing significant distinctions between various regions. These differences are particularly notable between the Tohoku and Tokai regions (*Z* = 6.3750, *p* < .001), the Tohoku and Sanyo regions (*Z* = 5.8520 *p* = .0000), the Kanto and Tokai regions (*Z* = 4.8797, *p* < .001), the Kanto and Sanyo regions (*Z* = 4.1804, *p* = .0008), the Hokuriku and Tokai regions (*Z* = 3.7734, *p* = 0.0040), and the Hokuriku and Sanyo regions (*Z* = 3.8285, *p* = .0032) in PC2. Additionally, there are significant differences in PC4 between the Kanto and Shikoku regions (*Z* = 3.3020, *p* = .0215) and the Shikoku and Kyushu regions (*Z* = 3.0875, *p* = .0423). It should be noted that there are no significant differences observed in PC1 and PC3, and the regional differences remain relatively limited. Furthermore, there are few sexual differences (see Supplementary S2), with some exceptions of PC3 and PC4 in the early phase of the Tokai region and PC4 in the middle phase of the Kanto region, which is supported by statistical tests between the sexes in each PC score (*U* = 15926, *p* = .8128 in PC1, *U* = 13925, *p* = .0690 in PC2, *U* = 12693, *p* = .0020 in PC3, and *U* = 13695, *p* = 0.040 in PC4 comparisons).

**Figure 5:**
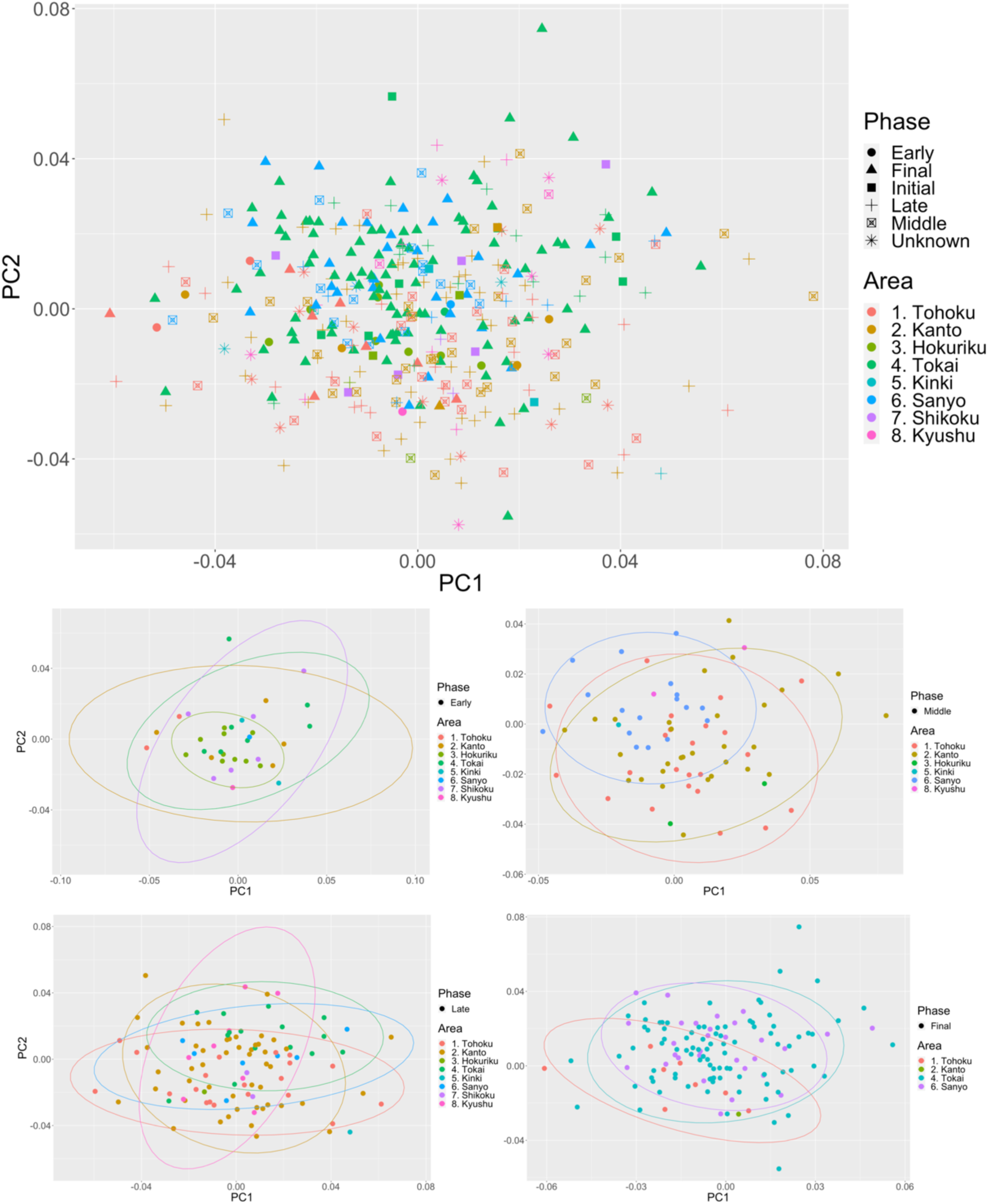
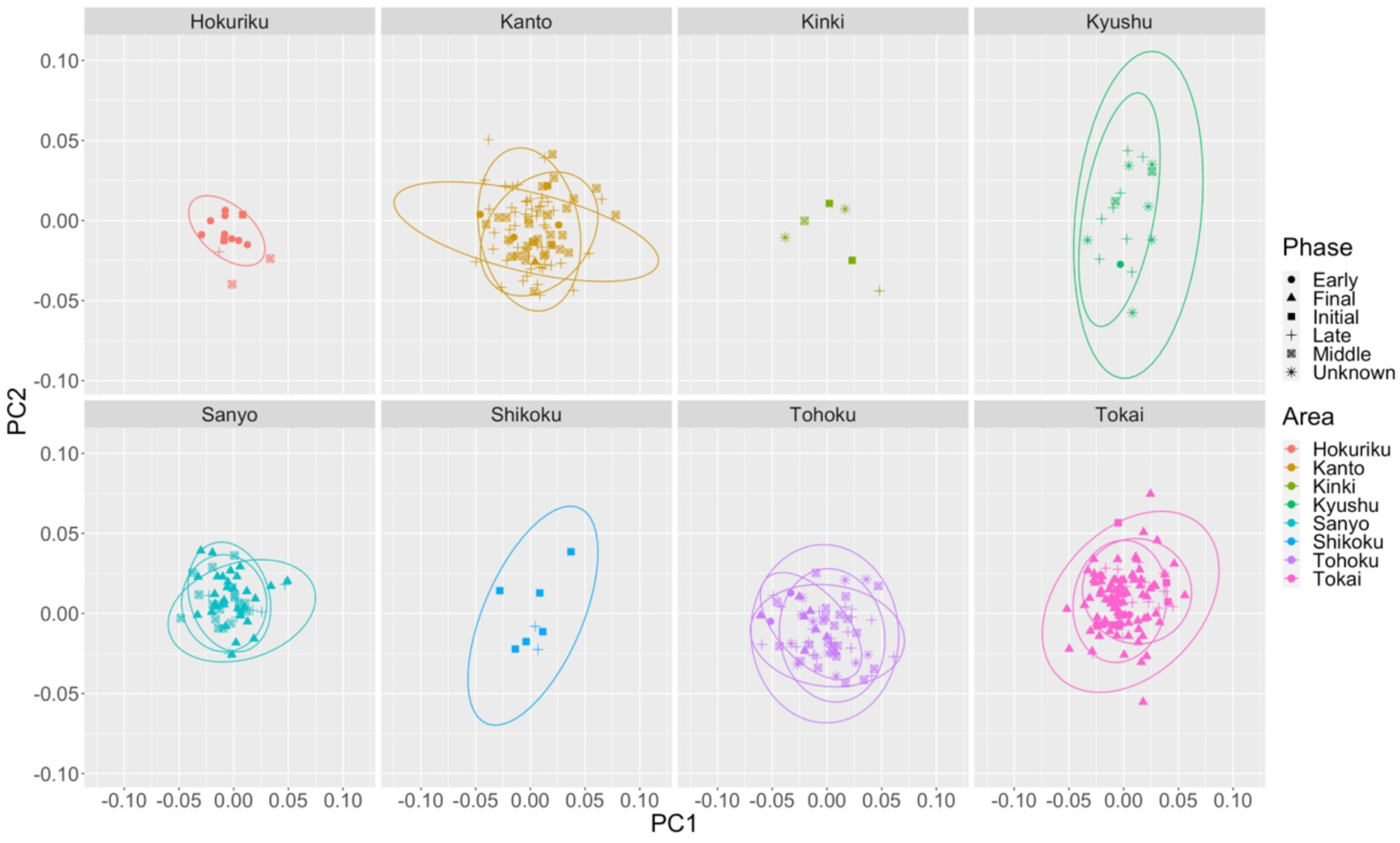
Principal component analysis (PCA) results for Jōmon crania.

**Figure 6:**
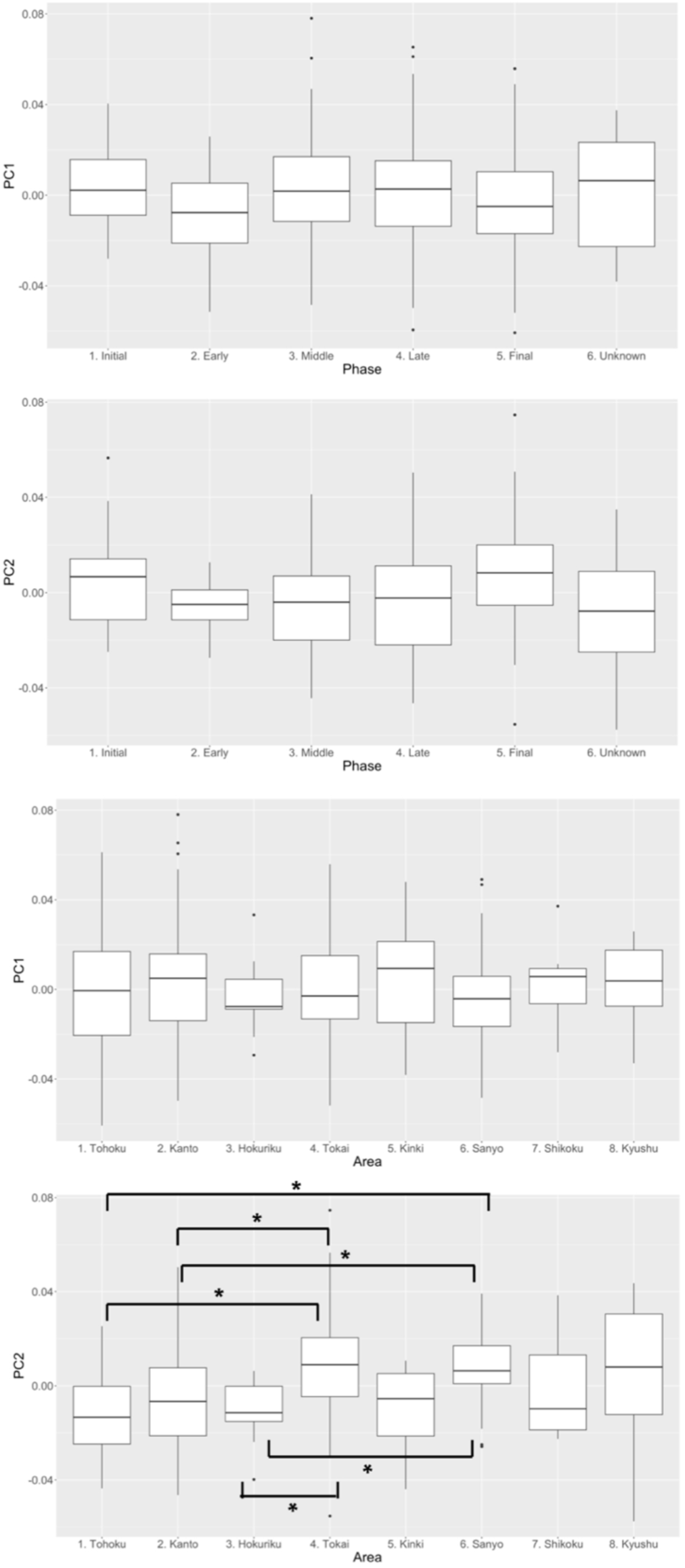
Boxplots of principal components (PC1 and PC2) by phase and by region (\**p* < .01)

The comparison results between the Jōmon and Yayoi populations are concisely summarized in Figure 7 and supplementary S3. To streamline the presentation, these figures show only PC1 and PC2, as all sites display substantial overlap in PC3 and PC4 (see supplementary S3). The most pronounced differences are observed in PC1, with one notable exception seen when comparing Nakazuma and Kuma–Nishioda sites, where the differences are more conspicuous in PC2. Figure 8 shows a visual representation of the configurational changes in landmarks for each comparison. These overall differences primarily relate to facial height and tooth position, specifically highlighting that individuals from Kuma–Nishioda site tend to exhibit taller facial features and a more anterior tooth placement.

**Figure 7:**
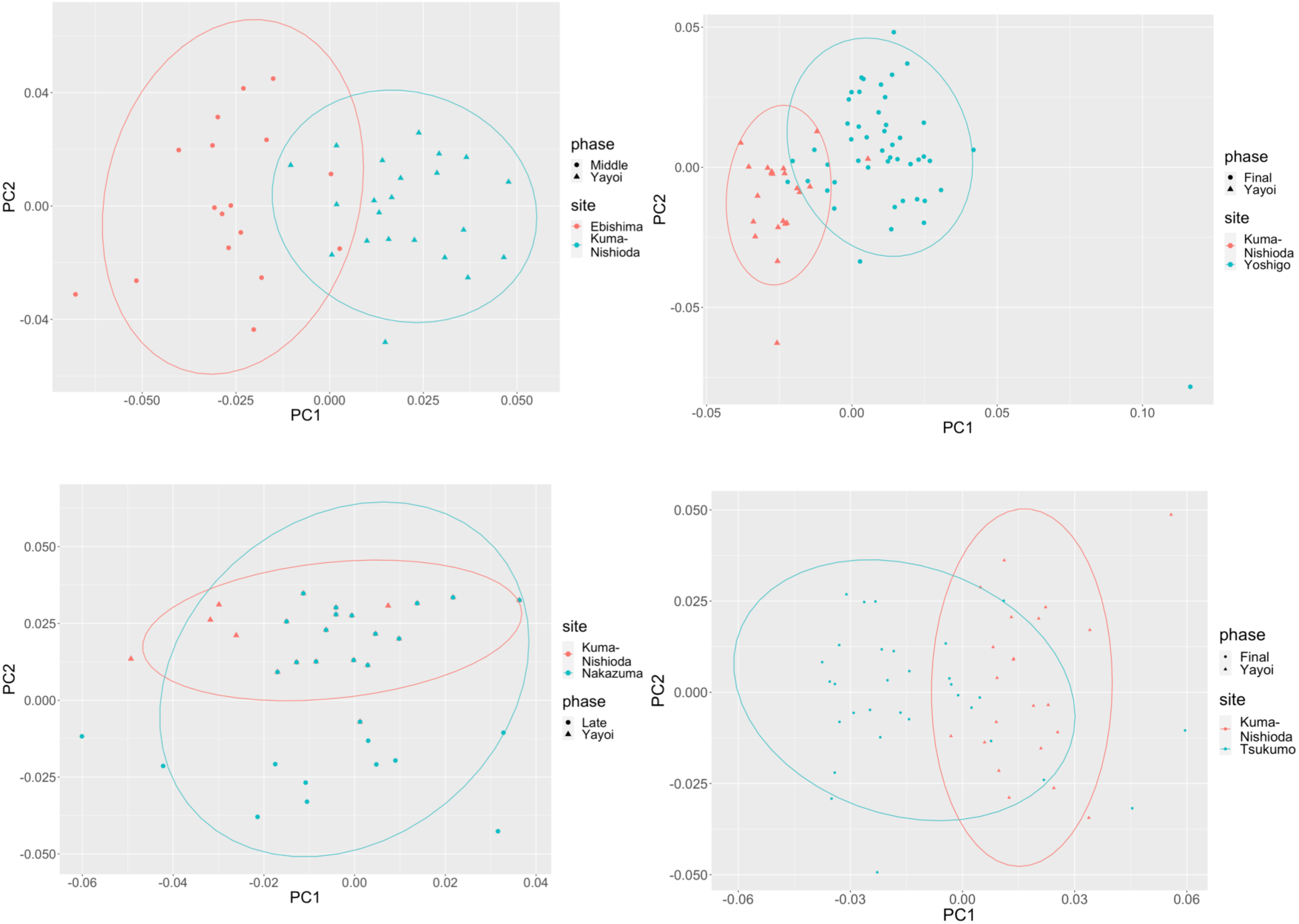
Comparative results between Jōmon sites (Ebishima, Nakazuma, Yoshigo, and Tsukumo) and Yayoi sites (Kuma–Nishioda). Refer to supplementary materials for results between each Jōmon site.

**Figure 8:**
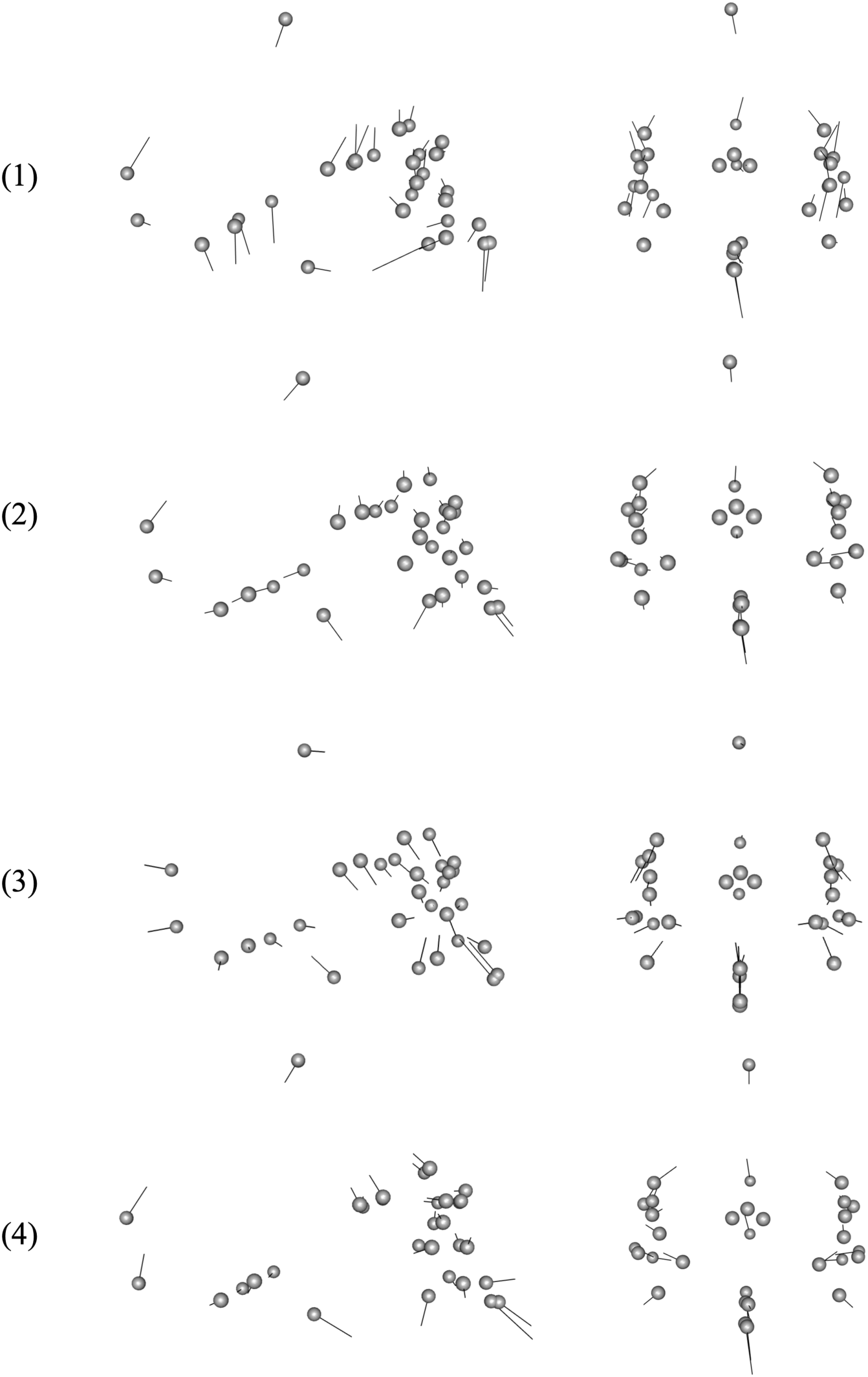
Landmark configuration changes in comparisons between some Jōmon sites and the Kuma–Nishioda site. Comparisons with the Kuma–Nishioda site for (1) Ebishima (middle Jōmon), (2) Nakazuma (late Jōmon), (3) Tsukumo (final Jōmon), and (4) Yoshigo (final Jōmon)

## Discussion

The PCA results presented above reveal that inter-phase differences among Jōmon skeletal remains are remarkably small, contrasting with some previous studies claiming inter-phase variation (Ogata 1981; Yamaguchi 1982b). Moreover, the findings suggest that inter-region differences are less pronounced than previously proposed by traditional biometric studies, which often suggest geographical clines from north to south (Fukase 2012; Kondo 2018; Yamaguchi 1982a). Especially, the boxplots of PCs in Figure 6 by region, with the exception of PC2, do not exhibit such clear clinical patterns (Figure 6). This discrepancy may be due to methodological differences between traditional biometrics and geometric morphometrics. As mentioned in the Introduction, geometric morphometrics could examine morphological variation as a whole, whereas traditional biometrics has to compare each measured distance independently. In-depth anthropological and anatomical investigations that encompass variations in other aspects of human skeletal remains may be required for a more comprehensive understanding of these phenomena.

Variations within phases and regions are more salient, and which align with the possibility that interactions between populations were continuous and widespread within Jōmon society. The comparative findings between Jōmon and Yayoi populations provide further support for the possibility, emphasizing continuous and extensive interactions between Jōmon communities. These results indicate that variations among Jōmon individuals from the four sites are relatively similar and different from, as well as more diverse than, the Yayoi population at Kuma–Nishioda site. It has been widely proposed that Yayoi populations, particularly in the northern Kyushu, are descended from migratory groups originating from the Korean peninsula (Naito 1981; Nakahashi & Nagai 1989; Nakao et al. 2023, 2024). If such a genetic bottleneck, resulting from this migration, indeed contributes to the reduced variation observed in the Yayoi people of the northern Kyushu, it suggests that the morphological and genetic diversity among the Jōmon populations was not relatively limited, but homogeneous across regions and phases. The comparison results of the Jōmon populations with those from other periods, including the Kofun period, also suggest that the morphological variations between the Jōmon and Kofun populations were not significantly different (Nakao et al. 2023, 2024).

Examining population interactions in the Jōmon period could provide important insights into their resilience to environmental fluctuations at the time. The Japanese archipelago during the Jōmon period experienced severe climatic changes, including the Younger Dryas, than the subsequent Yayoi (800–250 AD) and Kofun (250–600 AD) societies (Habu 2004; Imamura 1996; Matsugi 2011; Mizoguchi 2013; Tsude 2005). Many have claimed that prehistoric societies undergoing radical environmental fluctuations tended to be accompanied by societal change (e.g., Burroughs 2005; Maher, et al. 2011; Munoz et al. 2010; Schug 2020; Schug et al. 2023). The Jōmon society may be no exception. Relevant research has suggested that climate change has affected population size in many regions (e.g., Kodama 2003, Koyama 1984; Yasuda 1989), which may have influenced other aspects of the Jōmon society. However, reactions such as the emergence of social hierarchies, shifts in subsistence patterns, and significant technological developments that frequently appeared in other periods are conspicuously absent from the Jōmon society. Jōmon people predominantly adhered to a hunter-gatherer lifestyle and persisted in using stone tools for over ten millennia. While recent findings at Usu-Moshiri site in Hokkaido have revealed evidence of violent injuries in more than five individuals, such events remained relatively rare within the broader context of Jōmon period. This stands in stark contrast to the subsequent Yayoi period (Nakao et al. 2016, 2020; Nakagawa & Nakao 2017; Nakagawa et al. 2017a, 2017b, 2021; Mizoguchi 2013) and can be distinguished from patterns observed in other relevant regions worldwide (Frayer 1997; Lahr et al. 2016; Wendorf 1968). While the notion of a “static view” of the Jōmon society has been challenged over the years (Habu 2004; Kosugi et al. 2010), it remains clear that societal changes during this period were relatively less drastic, even in the face of more drastic climatic fluctuations.

One potential strategy for mitigating the impacts of external environmental stresses is sustained interaction between different populations. Anthropological evidence suggests that Jōmon people behaviorally adapted to the environmental fluctuations rather than physically (Buck et al. 2019). The continuous exchange of biological resources, knowledge, skills, and resources can also enhance the resilience and adaptability of populations in the face of environmental disruptions. Many relevant studies of genetic diversity in organisms other than humans support this possibility, including plants, various animal species, and whole ecosystems (Booy et al. 2000; Hughes and Stachowicz 2004; Hughes et al. 2008).

Archaeological evidence is also consistent with our results and interpretation. As mentioned in the Introduction, various types of Jōmon pottery (e.g., the Ento-kaso type in the early phase and the Funamoto type in the middle phase) were found in broader regions, and some types such as the Horinouchi type I and the Nakatsu type in the late phase were also distributed to distant regions and influenced the local type of pottery (i.e., the Shomyoji type). Although we admit that such a broad distribution of various types of pottery might be interpreted solely in terms of cultural transmission without any demic diffusion, by referring to the broader distribution pattern of pottery and lithics in the initial phase, some research has argued that the nomadic population in the Tokai region made an expedition to the southern Tohoku region and temporarily co-lived together with the local population (Abe 2010).

Even if the present results suggest that Jōmon people moved or interacted widely, it is still possible that such movements or interactions were not *continuous* but *discontinuous* with some drastic environmental changes such as the volcanic explosion and the 4.2ka event (Kajita et al. 2023; Kawahata 2019; Maeno and Taniguchi 2017). The frequency and degree of interactions naturally depend on the region and phase, as suggested by research on pottery distributions (e.g., Imamura 2011; Yano 2016). It should be noted, however, that certain regional interactions should be maintained to sustain homogeneity across different regions. How frequent and continuous interactions could maintain the morphological variations revealed in the present study should be explored in future work.

Some results from genetic or isotopic research on Jōmon skeletal remains, suggesting that Jōmon people interacted with populations from different regions, are consistent with our interpretation (Kawanishi 2023; Kusaka et al. 2021). Cooke et al. (2021) argued that Jōmon people in the mainland of the Japanese archipelago were isolated from other islands, which might have contributed to the morphological homogeneity of Jōmon people. Nevertheless, to strengthen or verify this hypothesis, further genetic data, and morphological data from other relevant regions overseas should be collected and combined (Adachi et al. 2021; Buck et al. 2024; Kanzawa-Kiriyama et al. 2019; Matsumura et al. 2021; Wang et al. 2021). For instance, our samples are much larger than the ones used in Buck et al. (2024), claiming that dietary factors influenced the shape of the Jōmon neurocranium, while our results should be also combined with isotopic dietary data in the future work.

While the dataset comprising larger sample sizes is acknowledged, potential sampling bias may exist, particularly due to restrictions on obtaining samples, notably in the northern Kyushu region. However, previous biometric studies have suggested that individuals from the northern Kyushu region do not exhibit significant differences compared to other regions (Yamaguchi 1982b). This contributes to the overall reliability of the findings presented in this study. Our study exclusively focuses on cranial 3D data, although previous investigations have examined other aspects of skeletal remains, such as limbs (Fukase et al. 1992; Mouri 1988), warranting a comprehensive examination of their 3D data.

The selection of landmarks in this study is a critical aspect that warrants consideration. While the chosen landmarks were selected for their clarity and suitability, the effectiveness of landmarks in capturing morphologically significant changes may vary depending on various factors such as geographic regions, time periods, age groups, and other contextual factors.

## Conclusion

The present study focused on the examination of prehistoric hunter-gatherer societies, specifically the Jōmon society. We employed geometric morphometric analysis on the 3D data of their crania to investigate their population interactions. Our findings revealed that the primary sources of variation were within phases and regions, rather than between them. This observation suggests that Jōmon people maintained extensive interactions across different populations and geographic regions. These sustained and widespread interactions might explain the relative stability of Jōmon society in comparison to other prehistoric periods in the Japanese archipelago and related regions.

## Ethical Statement

Ethical approval was not required for the present study according to the local legislation and institutional requirements (i.e., Japanese laws regulating archaeological human remains in Japan). Curators of all samples are described in supplementary S1.

## Author Contributions

All designed the research. HN, TN, and KN gathered the 3D data. HN and TK analyzed the data, and HN wrote the original draft. All edited it and approved the final manuscript.

## Acknowledgements

We thank for the following cultural property or archaeological centers, institutions, museums, and universities (see the detailed information for the supplementary material). Iwate Prefectural Museum, Osaki City Borad of Education, Ofunato City Museum, Kamikuroiwa Archaeological Museum, Shiura Historical Museum, Tohoku History Museum, Tome City History Museum, Niigata University of Health and Welfare, Yokohama City Archaeological Center, Hamamatsu City Museum, Matsudo City Borad of Education, Anjo City Museum of History, Okayama University of Science, Kasaoka City Borad of Education, Kyoto University, Takayama Village History Museum, National Museum of Nature and Science, Sakaki Town Archaeological Center, Sakiyama shell midden Museum, Shiga Prefectural Archaeological Center, Kagoshima Prefecture Archaeological Center, Toride City Archaeological Center, Shirokawa History Museum, Aomori Prefectural Museum, Chiba Prefectural Board of Education, Osaka Metropolitan University, Tahara Municipal Museum, The Tohoku University Museum, Minamichita Town Borad of Education, Toyama Prefecture Archaeological Center, Hirajo Public Hall, Nagoya City Archaeological Center, Nagoya City Museum, Kariya City History Museum, Kasukabe City History Museum, Mizuko shell midden Museum, Saitama City Urawa Museum, Kaizu History Museum, Tobinodai park Museum, Iwata City Borad of Education, and Kazuhiko Tanaka (Nagano Nishi high school). We also appreciate the following researchers for their highly valuable suggestions, comments, information, and cooperation: Kazuhiro Sakaue (National Museum of Science), Atsushi Fujisawa (The Tohoku University Museum), Takafumi Nara (Niigata University of Medicine and Welfare), Mikiko Abe (Osaka Metropolitan University), Naoto Tomioka (Okayama University of Science), and Ai Takeuchi (Nanzan University).

## Supplementary Materials

