## Supplementary figures and images for "Continuous and widespread population interactions in the Jōmon society via geometric morphometrics on 3D data of human crania"

### Supplementary S2

Plots of PCs in females

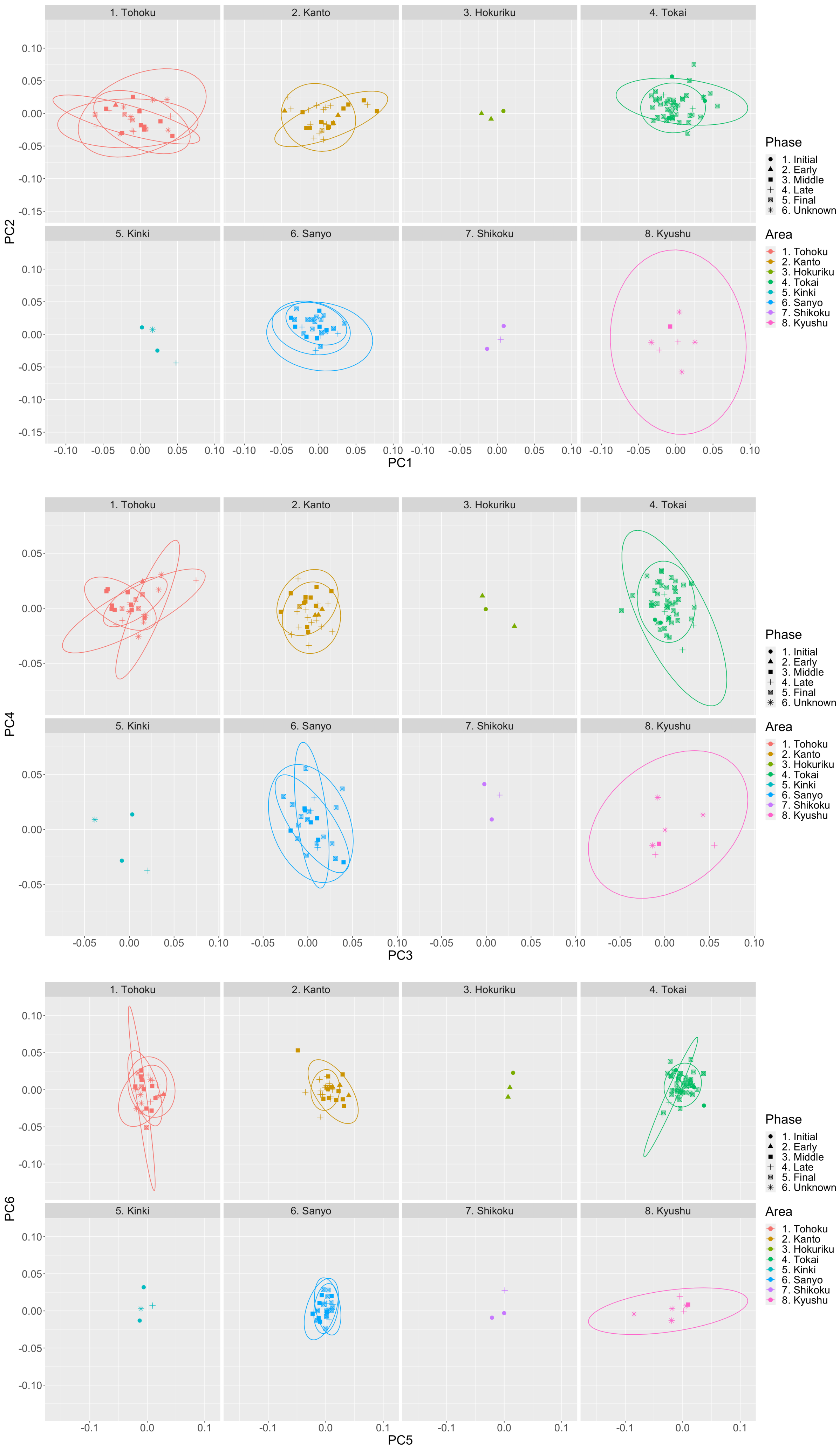

Plots of PCs in males

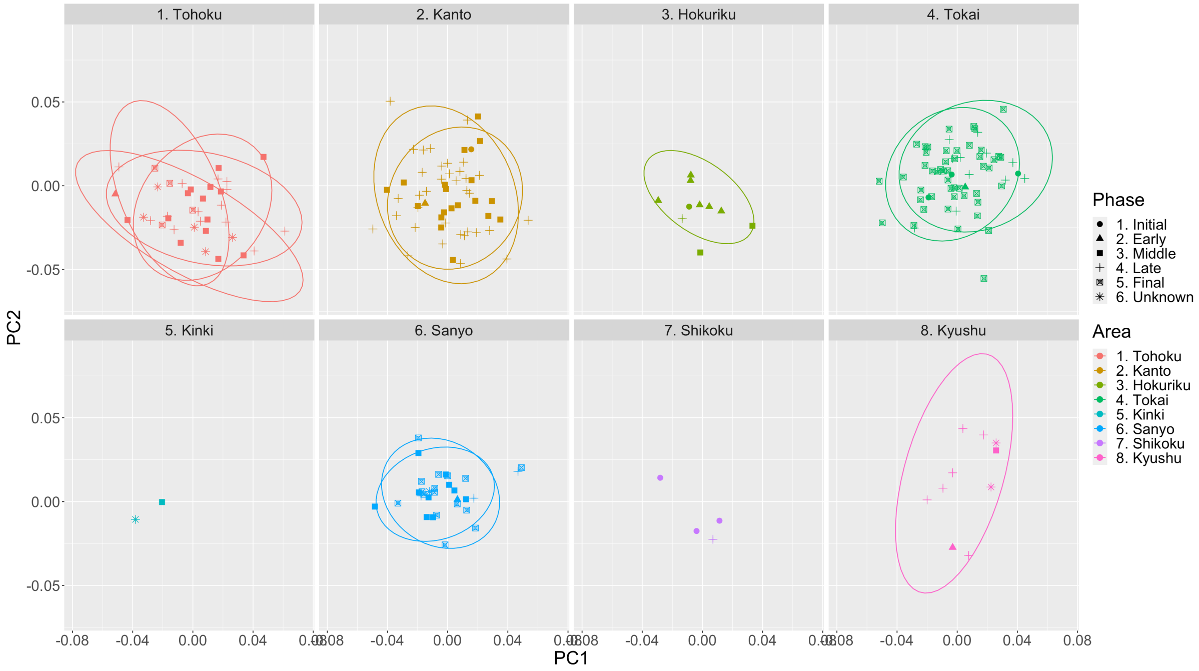


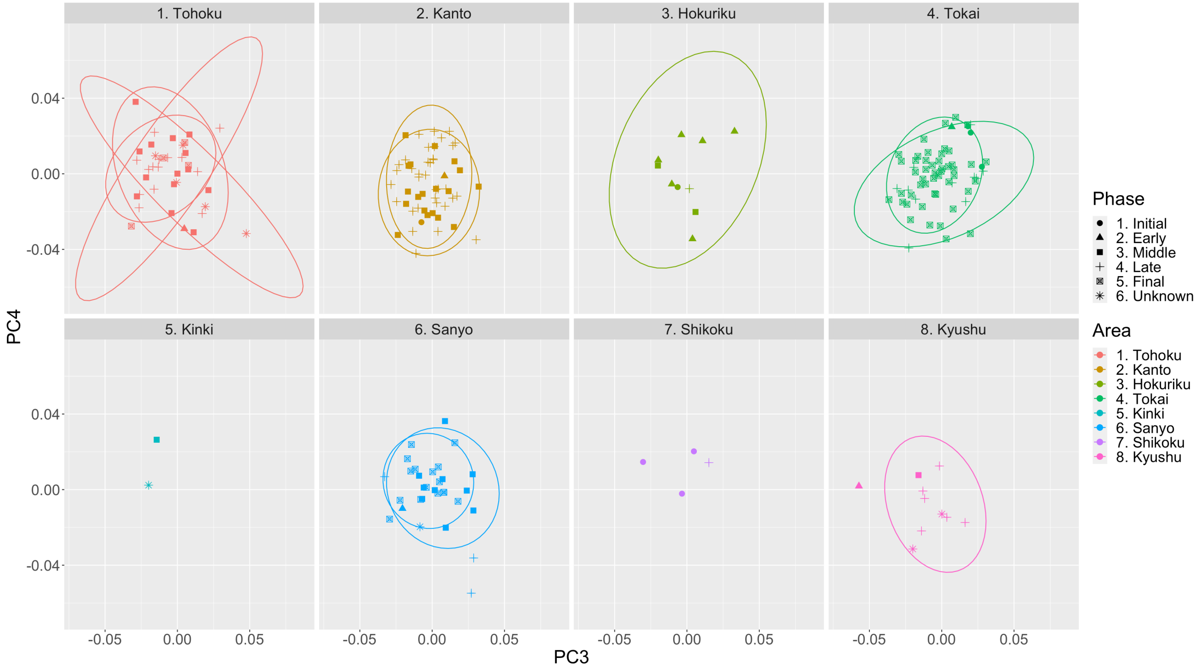


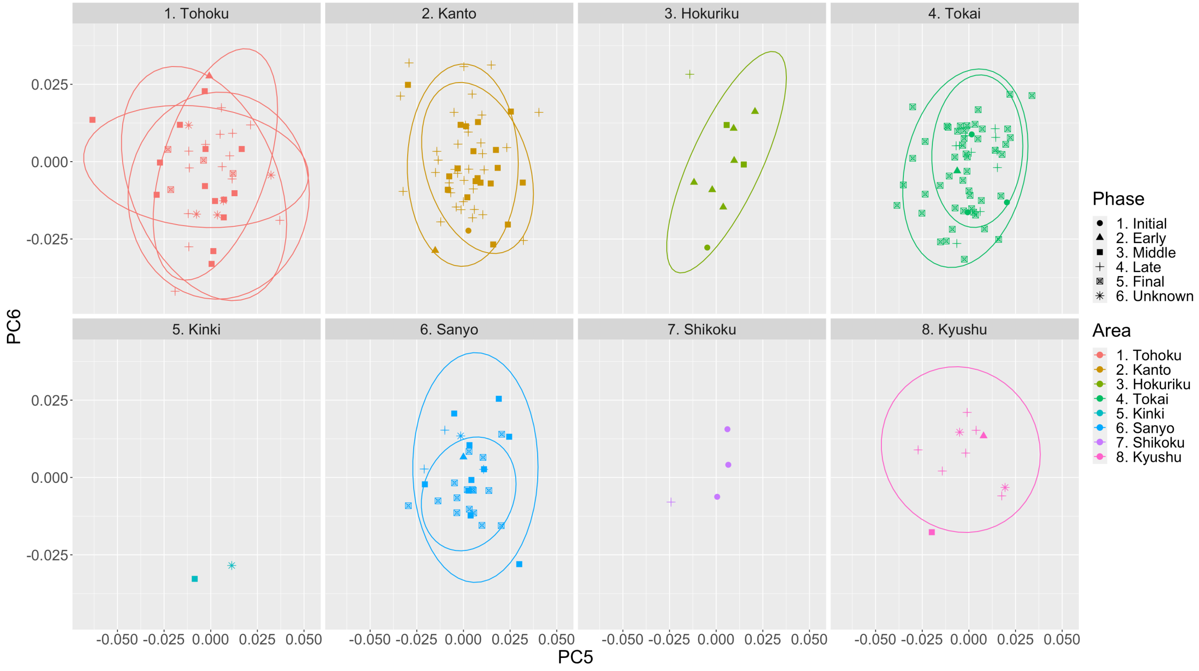

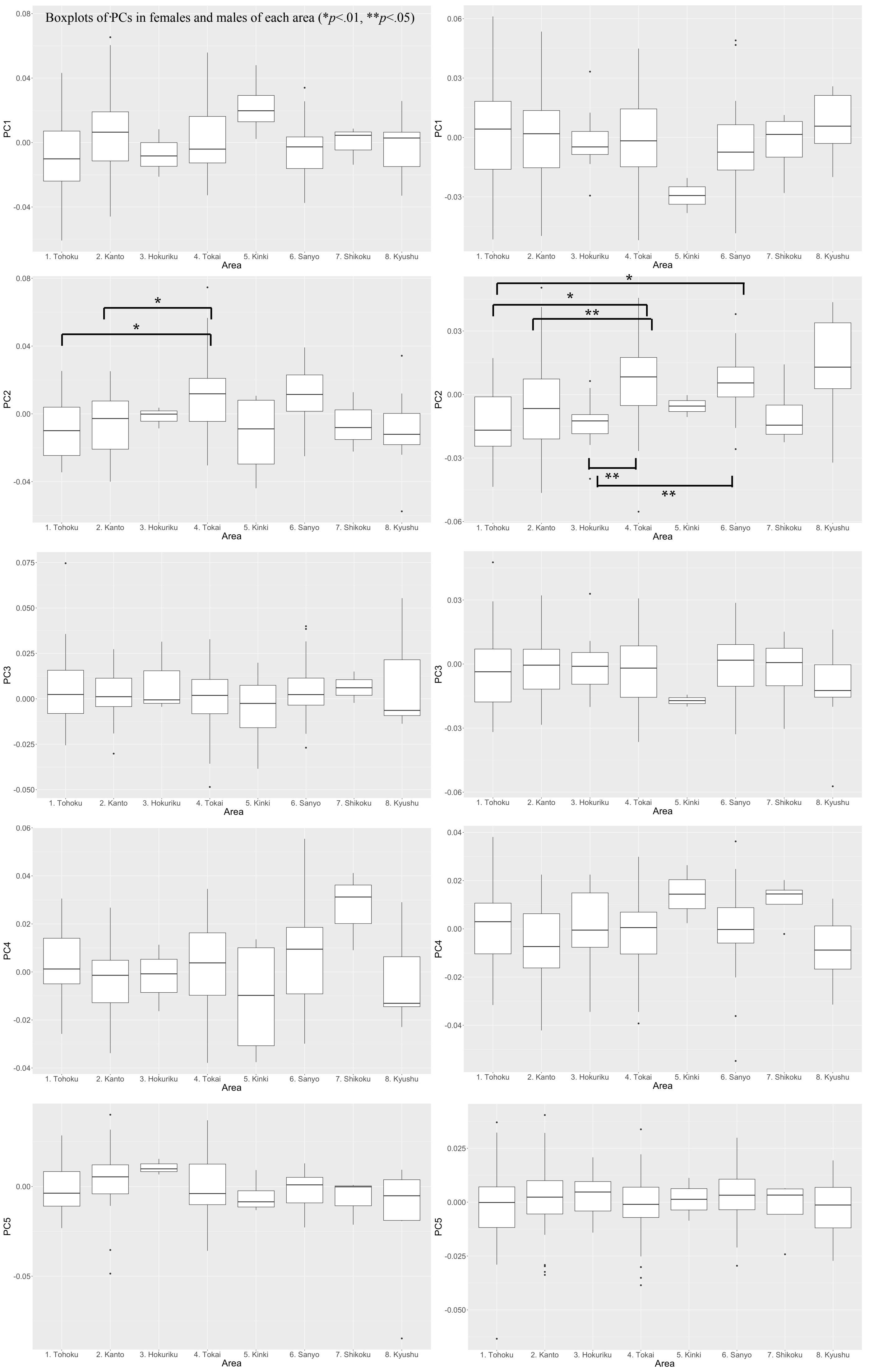

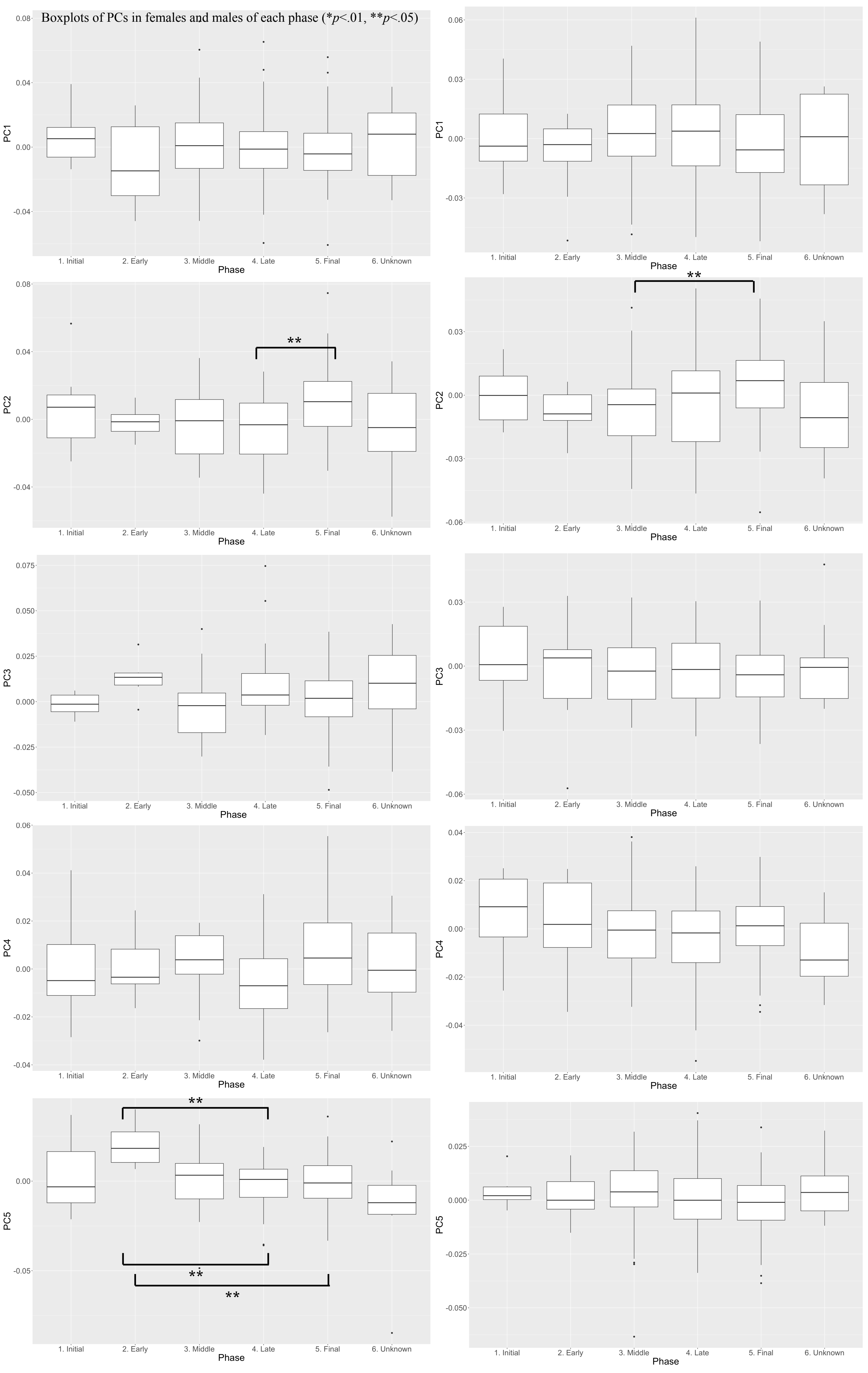
