## Supplementary S3 for "Continuous and widespread population interactions in the Jōmon society via geometric morphometrics on 3D data of human crania"

### PC1 and PC2 of comparisons between the Jomon sites

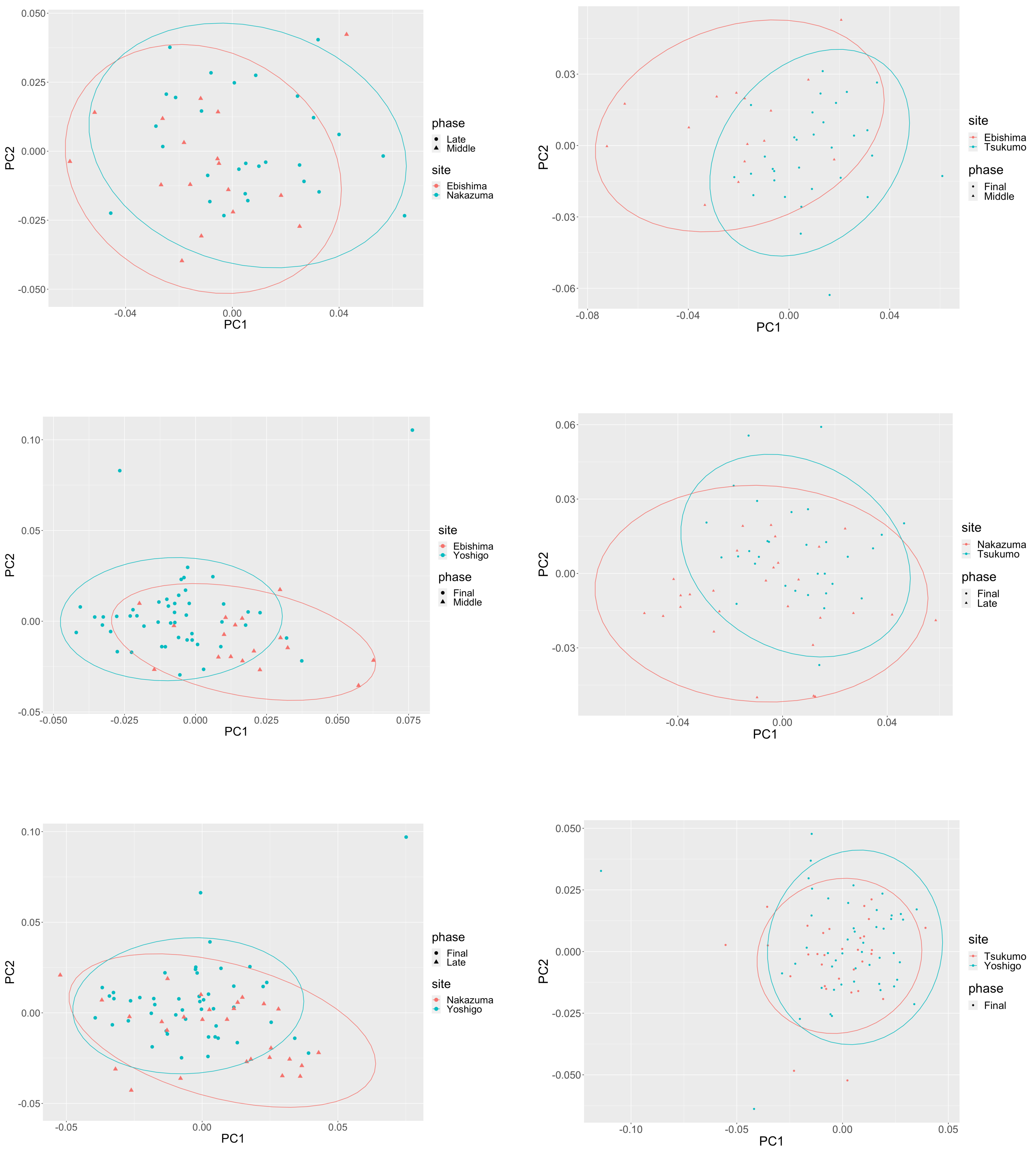

| Ebishima-Kuma | PC1 | PC2 | PC3 | PC4 | PC5 | PC6 | PC7 | PC8 | PC9 | PC10 | PC11 |
|---|---|---|---|---|---|---|---|---|---|---|---|
| Contribution rates | 0.184 | 0.109 | 0.095 | 0.073 | 0.071 | 0.053 | 0.046 | 0.040 | 0.035 | 0.032 | 0.028 |
| Cumulative proportion | 0.184 | 0.293 | 0.389 | 0.462 | 0.533 | 0.585 | 0.631 | 0.671 | 0.706 | 0.738 | 0.766 |

| Ebishima-nakazuma | PC1 | PC2 | PC3 | PC4 | PC5 | PC6 | PC7 | PC8 | PC9 | PC10 | PC11 | PC12 |
|---|---|---|---|---|---|---|---|---|---|---|---|---|
| Contribution rates | 0.172 | 0.093 | 0.086 | 0.081 | 0.066 | 0.056 | 0.051 | 0.040 | 0.036 | 0.031 | 0.026 | 0.024 |
| Cumulative proportion | 0.172 | 0.265 | 0.351 | 0.432 | 0.499 | 0.555 | 0.606 | 0.646 | 0.682 | 0.713 | 0.739 | 0.763 |

| Ebishima-tsukumo | PC1 | PC2 | PC3 | PC4 | PC5 | PC6 | PC7 | PC8 | PC9 | PC10 | PC11 | PC12 |
|---|---|---|---|---|---|---|---|---|---|---|---|---|
| Contribution rates | 0.161 | 0.107 | 0.089 | 0.075 | 0.065 | 0.056 | 0.049 | 0.039 | 0.038 | 0.030 | 0.030 | 0.025 |
| Cumulative proportion | 0.161 | 0.268 | 0.357 | 0.433 | 0.498 | 0.554 | 0.603 | 0.641 | 0.679 | 0.709 | 0.739 | 0.764 |

| Ebishima-yoshigp | PC1 | PC2 | PC3 | PC4 | PC5 | PC6 | PC7 | PC8 | PC9 | PC10 | PC11 | PC12 | PC13 |
|---|---|---|---|---|---|---|---|---|---|---|---|---|---|
| Contribution rates | 0.132 | 0.114 | 0.081 | 0.078 | 0.065 | 0.058 | 0.043 | 0.042 | 0.039 | 0.037 | 0.029 | 0.024 | 0.021 |
| Cumulative proportion | 0.132 | 0.246 | 0.327 | 0.405 | 0.470 | 0.528 | 0.571 | 0.613 | 0.652 | 0.689 | 0.718 | 0.743 | 0.763 |

| Kuma-nakazuma | PC1 | PC2 | PC3 | PC4 | PC5 | PC6 | PC7 | PC8 | PC9 | PC10 | PC11 | PC12 |
|---|---|---|---|---|---|---|---|---|---|---|---|---|
| Contribution rates | 0.130 | 0.126 | 0.091 | 0.074 | 0.059 | 0.057 | 0.047 | 0.042 | 0.037 | 0.033 | 0.029 | 0.025 |
| Cumulative proportion | 0.130 | 0.256 | 0.347 | 0.421 | 0.481 | 0.538 | 0.585 | 0.627 | 0.664 | 0.696 | 0.725 | 0.751 |

| Kuma-tsukumo | PC1 | PC2 | PC3 | PC4 | PC5 | PC6 | PC7 | PC8 | PC9 | PC10 | PC11 | PC12 |
|---|---|---|---|---|---|---|---|---|---|---|---|---|
| Contribution rates | 0.154 | 0.097 | 0.095 | 0.074 | 0.066 | 0.052 | 0.045 | 0.042 | 0.038 | 0.034 | 0.031 | 0.026 |
| Cumulative proportion | 0.154 | 0.251 | 0.345 | 0.419 | 0.485 | 0.537 | 0.583 | 0.624 | 0.662 | 0.696 | 0.727 | 0.754 |

| Kuma-yoshigo | PC1 | PC2 | PC3 | PC4 | PC5 | PC6 | PC7 | PC8 | PC9 | PC10 | PC11 | PC12 | PC13 | PC14 |
|---|---|---|---|---|---|---|---|---|---|---|---|---|---|---|
| Contribution rates | 0.128 | 0.098 | 0.093 | 0.078 | 0.062 | 0.052 | 0.046 | 0.043 | 0.038 | 0.032 | 0.028 | 0.025 | 0.024 | 0.020 |
| Cumulative proportion | 0.128 | 0.226 | 0.320 | 0.398 | 0.460 | 0.512 | 0.558 | 0.601 | 0.640 | 0.672 | 0.699 | 0.725 | 0.748 | 0.769 |

| Nakazuma-tsukumo | PC1 | PC2 | PC3 | PC4 | PC5 | PC6 | PC7 | PC8 | PC9 | PC10 | PC11 | PC12 | PC13 | PC14 |
|---|---|---|---|---|---|---|---|---|---|---|---|---|---|---|
| Contribution rates | 0.143 | 0.118 | 0.077 | 0.069 | 0.057 | 0.048 | 0.043 | 0.040 | 0.037 | 0.034 | 0.030 | 0.025 | 0.025 | 0.021 |
| Cumulative proportion | 0.143 | 0.261 | 0.338 | 0.407 | 0.464 | 0.512 | 0.555 | 0.596 | 0.633 | 0.667 | 0.697 | 0.722 | 0.747 | 0.769 |

| Yoshigo-nakazuma | PC1 | PC2 | PC3 | PC4 | PC5 | PC6 | PC7 | PC8 | PC9 | PC10 | PC11 | PC12 | PC13 | PC14 |
|---|---|---|---|---|---|---|---|---|---|---|---|---|---|---|
| Contribution rates | 0.126 | 0.115 | 0.078 | 0.070 | 0.056 | 0.051 | 0.045 | 0.041 | 0.037 | 0.035 | 0.030 | 0.027 | 0.024 | 0.021 |
| Cumulative proportion | 0.126 | 0.241 | 0.319 | 0.389 | 0.445 | 0.496 | 0.540 | 0.582 | 0.619 | 0.653 | 0.683 | 0.710 | 0.734 | 0.754 |

| Yoshigo-tsukumo | PC1 | PC2 | PC3 | PC4 | PC5 | PC6 | PC7 | PC8 | PC9 | PC10 | PC11 | PC12 | PC13 | PC14 | PC15 |
|---|---|---|---|---|---|---|---|---|---|---|---|---|---|---|---|
| Contribution rates | 0.128 | 0.088 | 0.084 | 0.073 | 0.062 | 0.050 | 0.049 | 0.040 | 0.037 | 0.036 | 0.029 | 0.026 | 0.024 | 0.022 | 0.019 |
| Cumulative proportion | 0.128 | 0.216 | 0.300 | 0.373 | 0.435 | 0.485 | 0.534 | 0.574 | 0.611 | 0.647 | 0.676 | 0.702 | 0.726 | 0.748 | 0.767 |

### PC3 and PC4 of comparisons between the Jomon and Yayoi sites

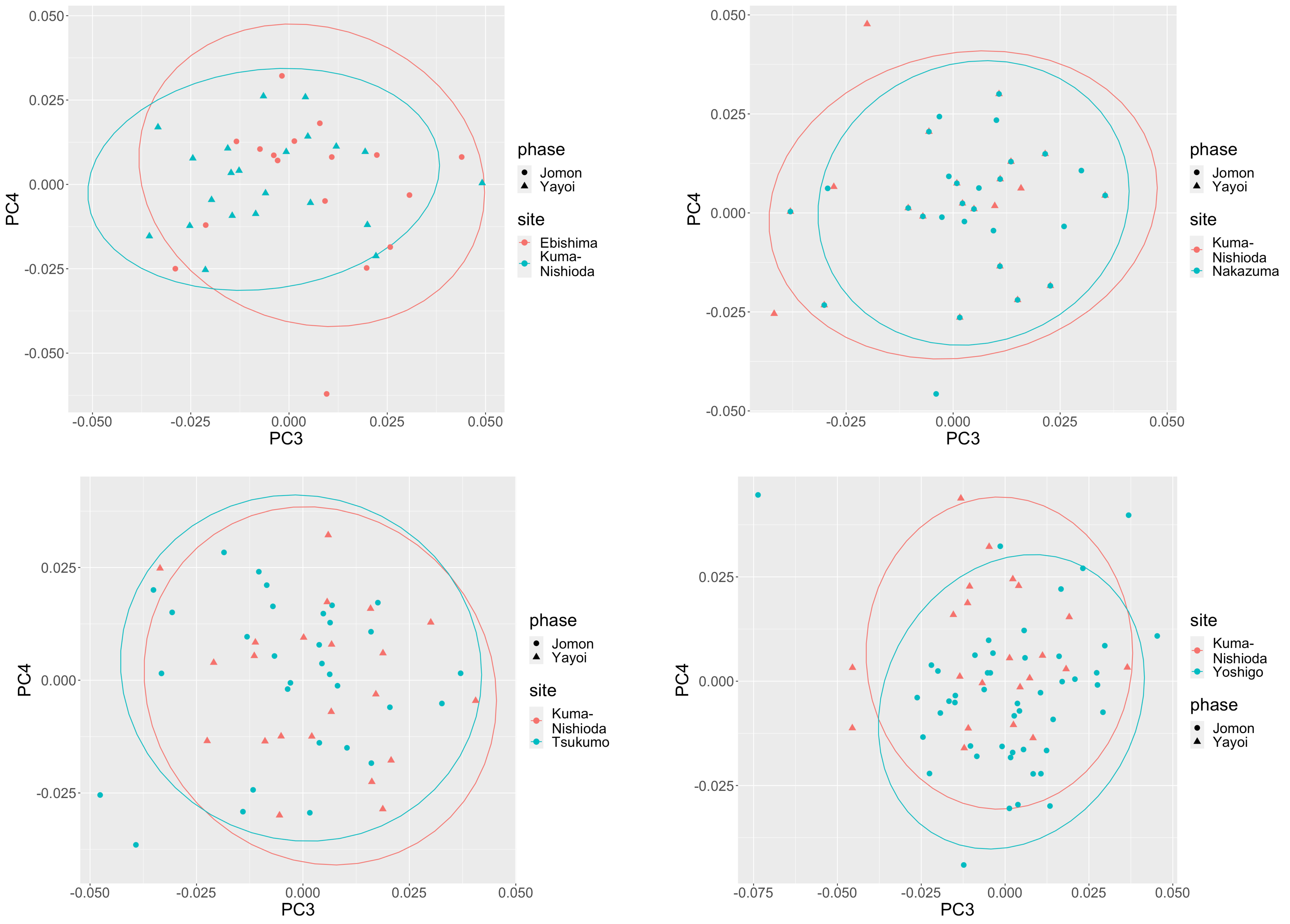
